# COVID-19 Detection on Chest X-Ray and CT Scan Images Using Multi-image Augmented Deep Learning Model

**DOI:** 10.1101/2020.07.15.205567

**Authors:** Kiran Purohit, Abhishek Kesarwani, Dakshina Ranjan Kisku, Mamata Dalui

**Affiliations:** Department of Computer Science and Engineering National Institute of Technology, Durgapur India

**Keywords:** Coronavirus, CNN, Image Augmentation, X-Ray images, CT Scan images

## Abstract

COVID-19 is posed as very infectious and deadly pneumonia type disease until recent time. Despite having lengthy testing time, RT-PCR is a proven testing methodology to detect coronavirus infection. Sometimes, it might give more false positive and false negative results than the desired rates. Therefore, to assist the traditional RT-PCR methodology for accurate clinical diagnosis, COVID-19 screening can be adopted with X-Ray and CT scan images of lung of an individual. This image based diagnosis will bring radical change in detecting coronavirus infection in human body with ease and having zero or near to zero false positives and false negatives rates. This paper reports a convolutional neural network (CNN) based multi-image augmentation technique for detecting COVID-19 in chest X-Ray and chest CT scan images of coronavirus suspected individuals. Multi-image augmentation makes use of discontinuity information obtained in the filtered images for increasing the number of effective examples for training the CNN model. With this approach, the proposed model exhibits higher classification accuracy around 95.38% and 98.97% for CT scan and X-Ray images respectively. CT scan images with multi-image augmentation achieves sensitivity of 94.78% and specificity of 95.98%, whereas X-Ray images with multi-image augmentation achieves sensitivity of 99.07% and specificity of 98.88%. Evaluation has been done on publicly available databases containing both chest X-Ray and CT scan images and the experimental results are also compared with ResNet-50 and VGG-16 models.

## 1. Introduction

Coronavirus disease or COVID-19 is an infectious disease which so far has infected millions of people and deaths are increasing day by day. Due to deadly infectious nature of coronavirus, it is spreading rapidly among people who are exposed to COVID-19 infected individuals

Due to unknown cause of pneumonia type infection and ability to generate new strain by mutation, it is almost impossible to have a cure in the form of vaccine or medicine for COVID-19 patients. In the affected countries, reverse transcription polymerase chain reaction or RT-PCR has been adopted as standard diagnostic method to detect viral nucleic acid as coronavirus infection in COVID-19 suspected individuals. The test takes 4-6 hours or even a whole day to give the results. As the test takes more time to generate the result compared to the time for spreading coronavirus among people and sometimes it gives false positive and false negative results. The false negative rate for SARS-CoV-2 RT-PCR testing is profoundly factor: most noteworthy in the initial 5 days after exposure (up to 67%), and least on day 8 after exposure (21%). In view of this investigation, the false negative rate for SARS-CoV-2 RT-PCR is very high, even at its least on day 8 post-exposure, or 3 days after symptoms. At its best, one out of five individuals associated with COVID-19 will test negative. RT-PCR tests showed 5% false positives rates. Therefore, to test the COVID-19 infection rapidly and in more efficient way, chest X-Ray or/and CT scan images of COVID-19 suspected individuals could be an answer. Moreover, time taken by RT-PCR test, false positive errors and shortage of test kits compared to coronavirus infected persons make it inefficient.

In contrast, X-Ray and CT scan images are widely accepted traditional form of diagnosing individuals for a number of diseases is a common practice adopted by radiologists and medics in healthcare and in medical imaging. The X-Ray and CT scan technologies have been used for several decades since its inception in medical diagnosis. In many highly affected regions or countries, it is difficult to provide sufficient number of RT-PCR test kits for testing COVID-19 infection for thousands of corona suspected people. Therefore, to address this issue, COVID detection can be made from chest X-Ray and CT scan images of corona suspected individuals who are suffering from COVID-19 symptoms.

The previous investigations uncover that the contaminated patients display distinct visual qualities in Chest X Ray pictures, as appeared in Figure 1. These qualities regularly incorporate multi‐focal, reciprocal ground‐glass opacities and inconsistent reticular (or reticulonodular) opacities in non-ICU patients, while thick aspiratory unions in ICU patients. Nonetheless, the manual investigation of these inconspicuous visual attributes on Chest X Ray pictures requires domain expertise and is challenging. Also, the exponential increment in the quantity of contaminated patients makes it hard to finish the finding in time, prompting high bleakness and mortality.

**Fig. 1:**
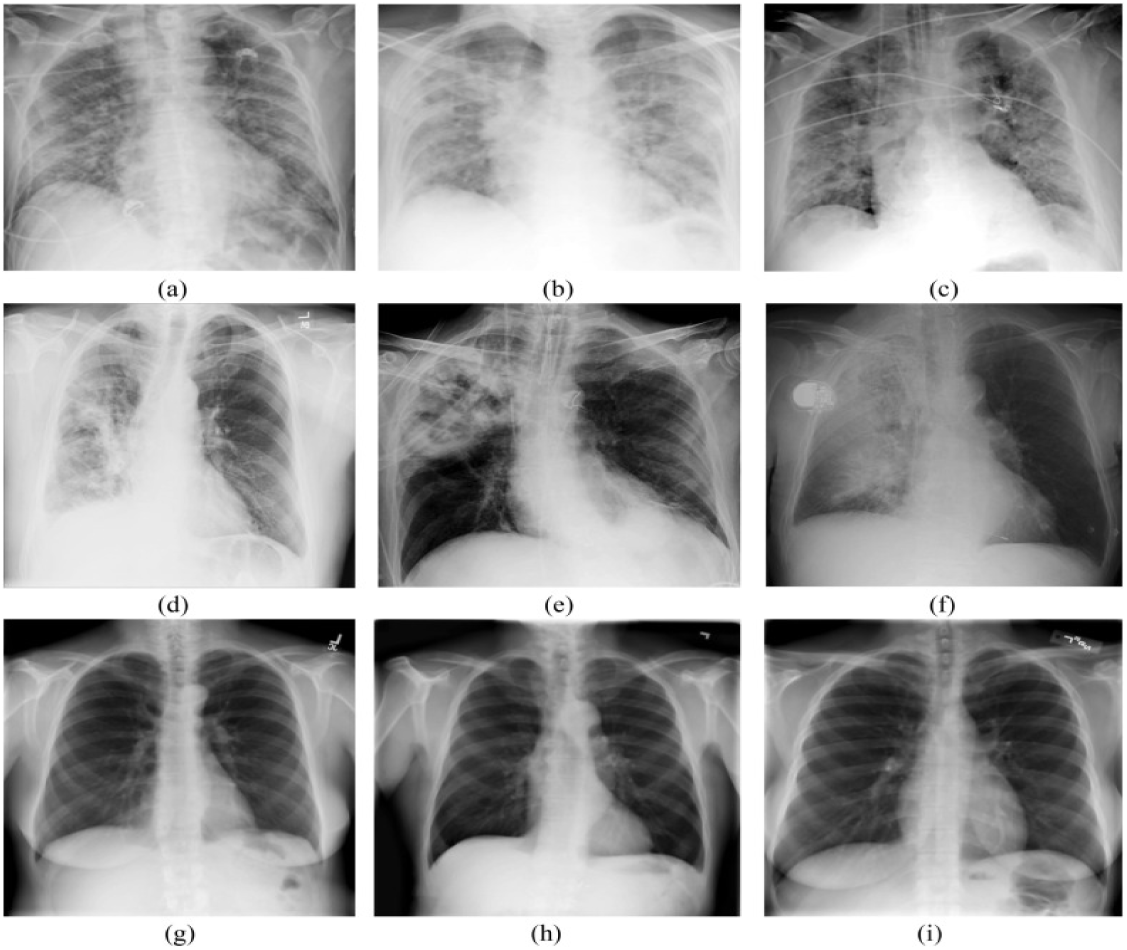
(a-c) COVID-19 infected chest X-Ray images, (d-f) Pneumonia infected chest X-Ray and (g-i) Normal chest X-Ray images [4].

For CT SCAN, imaging discoveries have mostly included single or numerous injuries, sketchy or segmental GGOs, and reticular markings that essentially followed peribronchovascular and subpleural disseminations. Interlobular septal thickening may be available, and pleural emission and extended mediastinal lymph hubs were infrequently observed.

To deal with the issues related to RT-PCR testing kit, we have come up with a solution by developing an AI-based application with image processing based multi-image augmentation technique for detecting COVID-19 infection in corona suspected persons. With this integrated framework, both X-Ray and CT scan images of chest can be tested for virus detection. This application makes use of multiple representations having sharp discontinuity information of same X-Ray and CT scan images, produced through first and second order derivative operators, are mixed up with visible band X-Ray and CT scan images for training the convolutional neural network (CNN) based deep learning model. This deep learning model has the ability to learn the underlying pattern of COVID-19 infected X-Ray and CT scan images in a more effective way from representative images as well as original images of the same person used for training. Moreover, with a simple configuration of CNN model, this work performs well for a range of COVID-19 infected X-Ray and CT scan images of chest.

The main objective of using deep learning model [19] is to achieve higher accuracy of classification with chest X-Ray and CT scan images by separating the COVID-19 cases from non-COVID-19 cases. It is well-known that to train a deep model, someone needs a large number of example images of both COVID-19 and non-COVID-19 individuals for making the learning of the model about the patterns more effective. To achieve this target, a number of representative images are generated using sharpening filters technique driven by first and second order derivative operators [10] and then these discontinuity information of the representations are mixed up with the original X-Ray and CT scan images separately and further, these large number of multi-image augmentation is used to train the CNN based deep learning model. The databases of X-Ray [6] and CT scan [24] images are publicly available in GitHub repository for the purpose of experiments. Both these datasets contain chest images of COVID-19 and non-COVID-19 individuals. The X-Ray database contains 67 COVID images and the same number of non-COVID images whereas CT scan database contains 345 COVID images and the same number of non-COVID images. To conduct the experiment, images are down sampled to 50×50 dimension from their original size. The random subsampling or holdout method is adopted to test the efficacy of the model. In holdout method, the whole dataset containing COVID positive and negative samples are divided into a number of ratios. It has been observed that when the number of training examples are increased, the model exhibits higher classification accuracy. Moreover, this result exhibits more consistency while layers are being changed in CNN based deep model. To evaluate the framework in a robust and effective way, a number of evaluation metrics such as classification accuracy, loss, area under ROC curve (AUC), precision, recall, F1 score and confusion matrix has been used. The values of these metrics have been determined on different ratios of training and test samples considering a number of layers in deep model. The model is correctly able to classify the chest X-Ray and CT scan images of COVID-19 cases from non-COVID-19 cases.

The paper is organized as follows. Section 2 introduces the proposed model to detect COVID-19 infections in CT Scan and X-Ray images of a chest. Section 3 presents experimental results, analysis and comparison among various deep learning models. Concluding remarks are made in the last section.

## 2 Proposed Work

In order to detect COVID-19 in chest X-Ray and CT Scan images of a lung the proposed work uses a number of sharpening filters such as Sobel, Prewitt, Roberts, Scharr, Laplacian, Canny, and a novel filter Hybrid.

### 2.1. Hybrid filter generation

To strengthen the process of detecting the sharp discontinuity on X-ray and CT Scan images, a novel edge detector is used, which is called Hybrid filter [1]. It uses both Canny [10] and Sobel [10] detector to normalize the noise content as well as provides the high frequency spatial information. This rare combination makes the operator very much useful for edge detection as well as image segmentation operations while texture properties of the image are enhanced. In this process, texton image [11] is used for unique texture which is generated from derivative of Gaussian [23]. Then the gradient of texton image is determined with the help of paired disk masks. The paired disk masks are the pair of half disk binary images. To get the texton gradient image, the pair of mask is convolved with the texton image and the distance is calculated between two convolved images with the use of Chi-square metric [13]. The output is obtained by the combination of Canny operator, Sobel operator and texton gradient image as given in Equation 1.

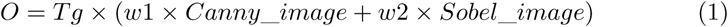

where, *O* is the output image, *T g* is the texton gradient image and *ω*1and *ω*2 are weights (*ω*1+ *ω*2 = 1). The design process of the Hybrid filter is shown in Figure 2.

### 2.2. Multi-image representation

Sharpening of an image [10] expands the contrast between dark and bright areas to draw out the features. Sharpening method uses a high pass kernel to an image. Sharpening is only inverse to the blurring. We decrease the edge content in case of blurring, and in sharpening, we increment the edge content. For noisy images, firstly smoothing is done and then a sharpening filter is applied.

**Fig. 2:**
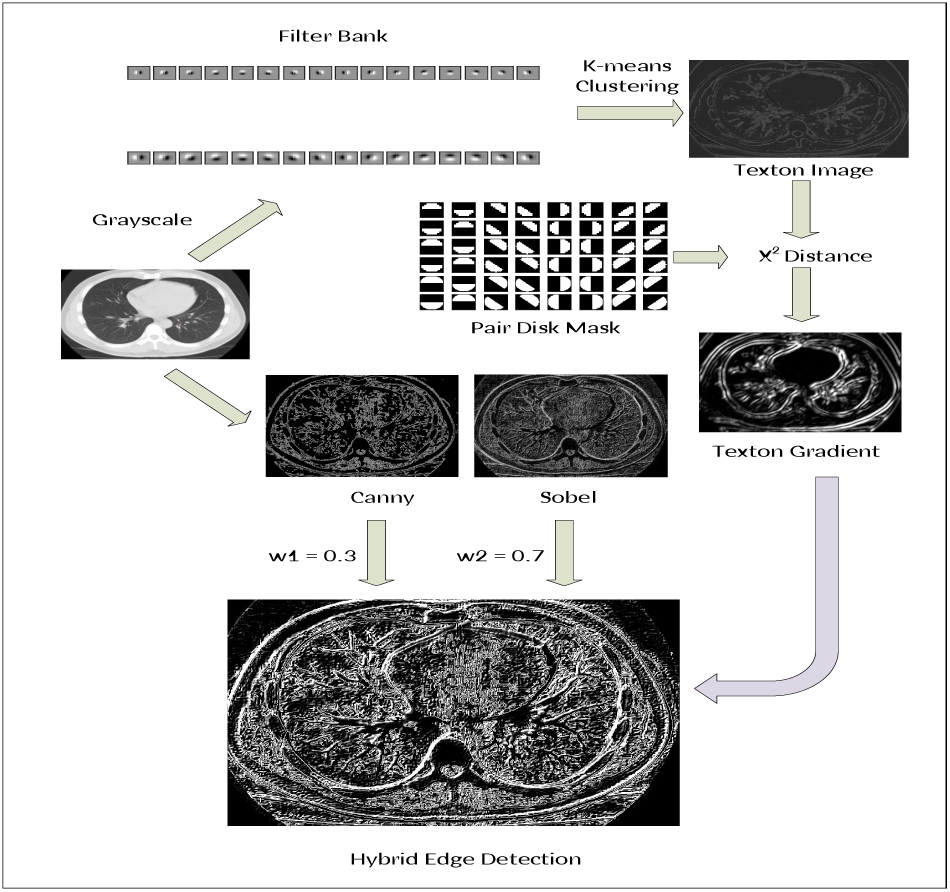
Design Process of the Hybrid Edge Detection Operator

The majority of the information about the shape of an image is enclosed in edges. Firstly, edges are distinguished in an image by the use of first and second-order derivative operators [10] and afterwards by enhancing those edges, image sharpness will increment and the image will become more clear. Consequently, detection of COVID-19 infection on X-Ray and CT Scan images will be more precise and accurate if we detect on the edged image.

The X-Ray [6] and CT scan [24] databases contain a smaller number of images which may not be useful for training the CNN model. Moreover, with small number of examples, desired classification accuracy may not be achieved. So, to resolve this issue we apply multi-image augmentation by increasing the number of examples as well as the diversity of available characteristics with X-Ray and CT Scan images. For multi-image augmentation, the input image is converted into grayscale and Histogram Equalization is applied to correct the contrast of input grayscale image. To achieve better image representation with discontinuity information, a number of first and second order edge detection operators such as Sobel, Prewitt, Roberts, Scharr, Laplacian, Canny, and newly develop Hybrid are applied. Then, the results of edge detection operator are appended to our dataset.

Edge detection operators perform a crucial job in separating low-level features or discovering information of the lungs. An edge is a sharp discontinuity change over the boundaries of the grey levels. In chest X-Ray and CT scan images, edges represent the lung boundaries, which occurs due to the change in the grey levels at these lung boundaries. Edges are determined to filter out relatively less basic and smaller details, for improving the processing speed, bringing down the complexity without the loss of the necessary information. Here, significant data is retained and non-essential data is separated out. We get more exact and accurate outcomes if the processing is performed on these edged images. Thus detection of COVID-19 infection on Chest X-Ray and CT Scan images is found to be more precise if detected on the edged image.

The Figure 3(a) shows a chest X-Ray (top left) and its multiple representations obtained by applying sharpening filters, of a person not infected by COVID-19 whereas the Figure 3(b) shows a chest X-Ray and its multiple representations obtained by applying sharpening filters, of COVID-19 infected person. Figure 3(c) shows a chest CT scan image and its multiple representations obtained by applying sharpening filters, of a person not infected by COVID-19 whereas Figure 3(d) shows a chest CT scan image and its multiple representations obtained by applying sharpening filters, of COVID-19 infected person.

**Fig. 3:**
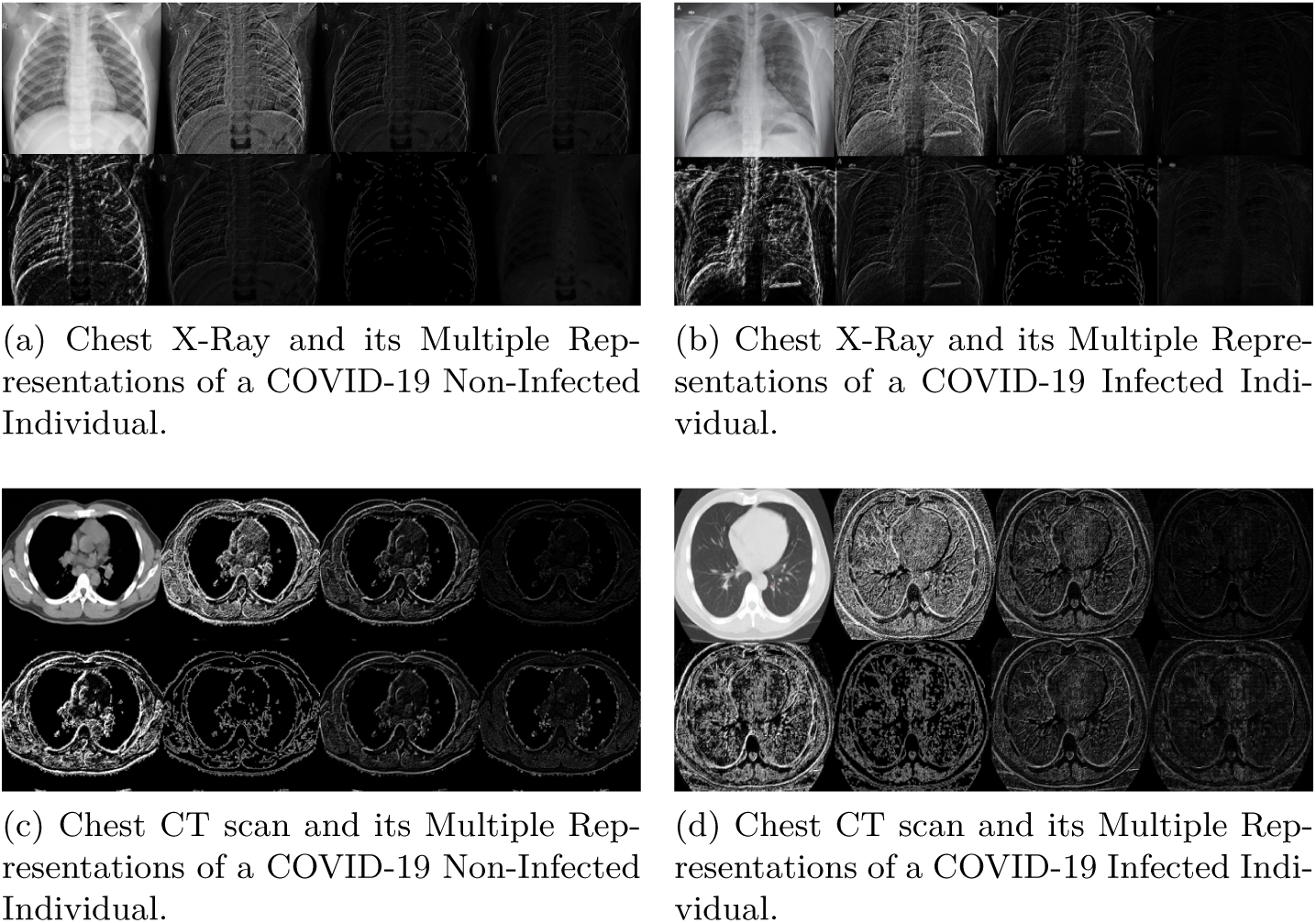
Multiple Representations of Chest X-Ray and CT scan images

### 2.3. Training and classification of CNN based deep learning model

To perform training and classification with a multi-image augmented CNN model the basic architecture of LeNet [7] model is exploited. It is used to predict COVID and non-COVID cases from CT Scan and X-ray images of lungs. The deep CNN model uses three layers such as convolutional, pooling and fully connected layers. Two activation functions viz. RELU and sigmoid are used. RELU is used after convolutional layer and sigmoid function is used for classification of test image into COVID and non-COVID classes. Figure 4 shows the deep learning model with a number of parameter. In the training stage, the standard stochastic gradient descent (SGD) optimizer with a batch size of 32 and a binary cross-entropy based loss function are used. The learning rate is set to 0.01, which is linearly decayed and the maximum epoch is set to 30. To conduct the experiment, images are down sampled to 50×50 dimension from their original size. The random subsampling or holdout method is adopted to test the efficacy of the model. In the holdout method, the whole dataset containing COVID positive and negative samples is divided into a number of ratios like 90:10, 80:20, 70:30 and 60:40 as training and testing samples. In order to alleviate overfitting of the model, multi-image augmentation is used for training the model using sharpening filters. This augmentation generates a large number of representative images carrying discontinuity information.

**Fig. 4:**
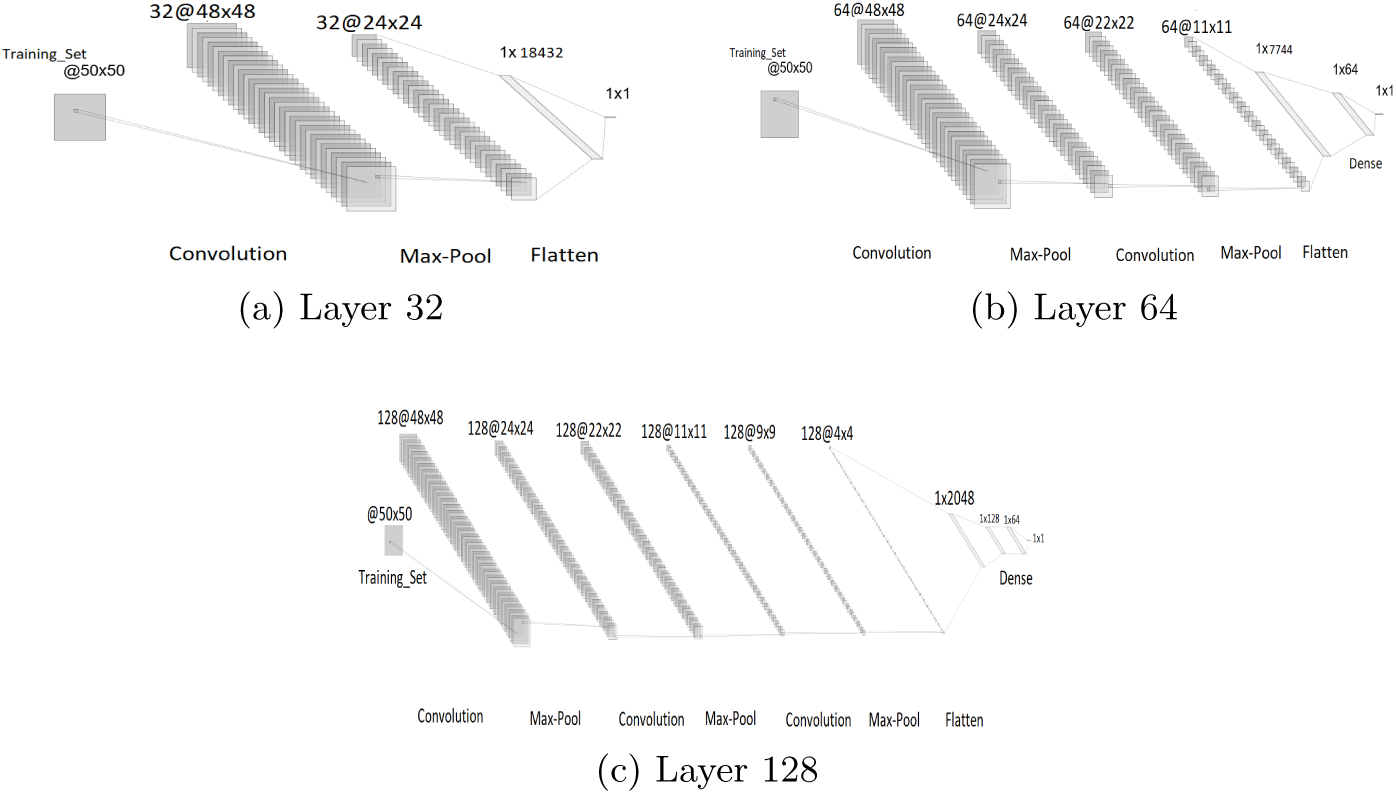
Convolution Neural Network of layer size 32, 64, and 128

Steps for detection of coronavirus infection in X-Ray and CT Scan images of the suspected individuals:

**Step 1:** Accept the coloured input images from the data set.

**Step 2:** Convert the image into grayscale.

**Step 3:** Down sample the images to 50×50 dimension from their original size.

**Step 4:** In CNN based deep learning model we choose *layer*_*sizes* = [32, 64, 128], *dense*_*layers* = [0, 1, 2], and *conv*_*layers* = [1, 2, 3].

**Step 5:** Convolution with a 3×3 filter size is applied.

**Step 6:** Activation function RELU is used after convolutional layer.

**Step 7:** Then Max Pooling is applied with 2×2 filter size.

**Step 8:** Go to Step 5 if *conv*_*layer* 1 *>* 0

**Step 9:** Flatten the matrix.

**Step 10:** Activation function Sigmoid is used for classification of test image into COVID and non-COVID classes.

**Step 11:** In training stage, the standard first-order stochastic gradient descent optimizer with a batch size of 32, maximum epochs 30 and binary cross entropy-based loss function are used.

**Step 12:** The random subsampling or holdout method is adopted. Whole dataset is divided into a number of ratios like 90:10, 80:20, 70:30 and 60:40 as training and testing samples.

**Step 13:** Various Evaluation Metrics are calculated such as classification accuracy, loss, area under ROC curve (AUC), precision, recall (Sensitivity), Specificity and F1 score.

**Step 14:** Once the model is trained, one can easily predict whether the individual is affected by Coronavirus infection or not.

## 3. Evaluation

### 3.1. Experimental protocol and metrics

The proposed multi-image augmented deep learning model, ResNet-50 and VGG-16 all are implemented using python code in Jupyter Notebook and Keras package with TensorFlow [8] on Intel(R) Core(TM) i5-8365U Processor with NVIDIA Geforce GTX 1050 graphical processing unit of 4GB and 16GB RAM. To evaluate the framework in a robust and effective way, a number of evaluation metrics such as classification accuracy, loss, area under ROC curve (AUC), precision, sensitivity, specificity and F1 score have been used. The values of these metrics have been determined on different ratios of training and test samples considering a number of layers in deep model. The model is correctly able to classify the chest X-Ray and CT scan images of COVID-19 cases from non-COVID-19 cases. The evaluation protocol makes use of holdout method where the whole dataset containing COVID positive and negative samples are divided into a number of ratios. During testing, the layers are changed in the augmented based deep learning model to realize the consistency of the model.

### 3.2. Experimental results

It has been observed that when the number of training examples are increased, the model exhibits higher classification accuracy. For traditional train and test ratio of 70:30 the classification accuracy is obtained around 95.38% and 98.97% for CT scan and X-Ray images respectively. The data is randomly splited into training and testing sets for evaluation. The experiment is conducted twice for every split and it is noted that the accuracy of the model comes similar on both evaluations. F1 scores remain slight similar and model loss (%) changes as the layer size is increased while setting the layer size to different values. For less training samples, a smaller layer size gives good results whereas if the training samples are increased then more layers are required for better accuracy. The proposed deep augmented model is also compared with ResNet-50 [2] and VGG-16 [17] architecture.

Visual Geometry Group(VGG) a research group from Oxford University in the United Kingdom proposed VGG networks. There are two commonly used networks, i.e., VGG-19 and VGG-16, here ’19’ represents 19 deep layers consists of 3 fully connected layers and 16 convolutional layers whereas the ’16’ means 3 fully connected layers and 13 convolutional layers. The size of input image in this network is 224× 224× 3. VGG network uses the pre-trained model on the ImageNet dataset which cannot improve the classification accuracy of COVID-19 screening as it contains a new set of images with labels. ResNet is known as ”residual network” which contains two core factors, i.e., width and depth of the neural network. These two core factors decide the complexity of the neural network. As the depth of a neural network increase, the training error increases. So, the residual network is used to solve this problem and increase the network performance (precision and accuracy) as compared to the traditional neural model. The commonly used residual networks are ResNet 18, 34 (2-deep layers) and ResNet 50, 101, 152 (3-deep layers). ResNet does not perform well in our case due to less amount of training datasets.

In this study, the proposed deep augmented model achieves the sensitivity of 99.07% when the ratio of train and test samples is 70:30 and specificity of 98.88% when the ratio of train and test samples is 70:30 and layer size 32 on the X-Ray dataset that contains 536 images of COVID-19 and 536 images non-COVID-19 including augmented images. The CT Scan dataset contains 2760 images of COVID-19 and 2760 images of non-COVID-19 individuals including augmented images. The sensitivity of 94.78% and specificity of 95.98% are achieved when ratio of train and test samples is 70:30 and layer size is 64. The results are shown in Table 1 and Table 2 for original and augmented images respectively. The model is able to correctly classify the chest X-Ray and CT scan images of COVID-19 cases from non-COVID-19 cases. Figure 5 shows the confusion matrix for the standard ratio of train and test samples i.e., 70:30 for both CT Scan and X-Ray images with multi-image augmentation and without multi-image augmentation on three different architectures as proposed. ROC curves for CT Scan and X-Ray images exhibiting higher accuracy when the ratio of train and test is 70:30 are shown in Figure 6.

**Table 1:** COVID-19 screening performance in Original images with CT Scan and X-Ray datasets

|  | CT Scan Images |  |  |  |  | X-Ray Images |  |  |  |  |
| --- | --- | --- | --- | --- | --- | --- | --- | --- | --- | --- |
|  | Proposed |  |  | Resnet | VGG | Proposed |  |  | Resnet | VGG |
| Train-Test | Layers |  |  |  |  | Layers |  |  |  |  |
| 60%-40% | 32 | 64 | 128 | 50 | 16 | 32 | 64 | 128 | 50 | 16 |
| Accuracy (%) | 88.7 | 90.87 | 87.68 | 88.7 | 78.7 | 98.51 | 99.25 | 98.51 | 50 | 98.51 |
| Loss (%) | 31.45 | 36.13 | 65.75 | 70.9 | 45.73 | 6.49 | 2.59 | 13.16 | 23.89 | 6.25 |
| AUC (%) | 94.5 | 95.2 | 93.5 | 88.7 | 78.7 | 99.9 | 100 | 98.5 | 50 | 98.5 |
| Precision | 0.86 | 0.9 | 0.89 | 0.91 | 0.72 | 0.97 | 1 | 0.97 | 0.5 | 0.97 |
| Sensitivity (%) | 92.17 | 92.17 | 85.51 | 85.5 | 93.62 | 100 | 98.51 | 100 | 1 | 100 |
| Specificity (%) | 85.22 | 89.57 | 89.86 | 91.88 | 63.76 | 97.01 | 100 | 97.01 | 0 | 97.01 |
| F1 Score | 0.89 | 0.91 | 0.87 | 0.88 | 0.81 | 0.99 | 0.99 | 0.99 | 0.67 | 0.99 |
| 70%-30% | 32 | 64 | 128 | 50 | 16 | 32 | 64 | 128 | 50 | 16 |
| Accuracy (%) | 89.57 | 92.03 | 90.29 | 78.84 | 77.54 | 99.25 | 97.1 | 100 | 50 | 98.51 |
| Loss (%) | 29.79 | 26.81 | 50.84 | 15.6 | 45 | 5.21 | 12.09 | 2.22 | 55.6 | 6.19 |
| AUC (%) | 94.8 | 95.9 | 95.7 | 78.8 | 77.5 | 100 | 99.8 | 100 | 50 | 98.5 |
| Precision | 0.91 | 0.93 | 0.87 | 0.71 | 0.76 | 0.99 | 1 | 1 | 0.5 | 0.97 |
| Sensitivity (%) | 87.25 | 91.01 | 95.07 | 97.39 | 81.15 | 100 | 94.03 | 100 | 88.05 | 100 |
| Specificity (%) | 91.88 | 93.04 | 85.51 | 60.28 | 73.91 | 98.51 | 100 | 100 | 11.94 | 97.01 |
| F1 Score | 0.89 | 0.92 | 0.91 | 0.82 | 0.78 | 0.99 | 0.97 | 1 | 0.64 | 0.99 |
| 80%-20% | 32 | 64 | 128 | 50 | 16 | 32 | 64 | 128 | 50 | 16 |
| Accuracy (%) | 94.06 | 95.22 | 94.35 | 93.48 | 78.7 | 99.25 | 97.01 | 99.25 | 51.49 | 98.51 |
| Loss (%) | 18.93 | 17.16 | 41.09 | 28.3 | 42.97 | 3.84 | 6.27 | 12.05 | 10.57 | 5.02 |
| AUC (%) | 98.1 | 98 | 96.8 | 93.5 | 78.7 | 100 | 100 | 98.6 | 51.5 | 98.5 |
| Precision | 0.93 | 0.95 | 0.94 | 0.93 | 0.72 | 0.99 | 1 | 0.99 | 1 | 0.97 |
| Sensitivity (%) | 95.07 | 95.65 | 94.78 | 94.49 | 93.04 | 100 | 94.03 | 100 | 2.98 | 100 |
| Specificity (%) | 93.04 | 94.78 | 93.91 | 92.46 | 64.34 | 98.51 | 100 | 98.51 | 1 | 97.01 |
| F1 Score | 0.94 | 0.95 | 0.94 | 0.94 | 0.81 | 0.99 | 0.97 | 0.99 | 0.06 | 0.99 |
| 90%-10% | 32 | 64 | 128 | 50 | 16 | 32 | 64 | 128 | 50 | 16 |
| Accuracy (%) | 94.93 | 98.26 | 98.26 | 94.49 | 83.33 | 99.25 | 100 | 99.25 | 50 | 99.25 |
| Loss (%) | 20.65 | 9.47 | 11.68 | 31.83 | 40.26 | 3.21 | 1.08 | 2.2 | 68.33 | 3.95 |
| AUC (%) | 98.4 | 99.5 | 99 | 0.91 | 83.3 | 100 | 100 | 100 | 50 | 99.3 |
| Precision | 0.97 | 0.97 | 0.98 | 0.91 | 0.79 | 0.99 | 1 | 0.99 | 0.5 | 0.99 |
| Sensitivity (%) | 92.46 | 99.42 | 98.55 | 98.26 | 91.01 | 100 | 100 | 100 | 1 | 100 |
| Specificity (%) | 97.39 | 97.1 | 97.97 | 90.72 | 75.65 | 98.51 | 100 | 98.51 | 0 | 98.5 |
| F1 Score | 0.95 | 0.98 | 0.98 | 0.95 | 0.85 | 0.99 | 1 | 0.99 | 0.67 | 0.99 |

**Table 2:** COVID-19 screening performance in Augmented images with CT Scan and X-Ray datasets

|  | CT Scan Images |  |  |  |  | X-Ray Images |  |  |  |  |
| --- | --- | --- | --- | --- | --- | --- | --- | --- | --- | --- |
|  | Proposed |  |  | Resnet | VGG | Proposed |  |  | Resnet | VGG |
| Train-Test | Layers |  |  |  |  | Layers |  |  |  |  |
| 60%-40% | 32 | 64 | 128 | 50 | 16 | 32 | 64 | 128 | 50 | 16 |
| Accuracy (%) | 93.46 | 92.54 | 91.59 | 85.53 | 85.07 | 98.88 | 98.79 | 98.04 | 93.66 | 90.39 |
| Loss (%) | 22.82 | 48.46 | 65.33 | 59.9 | 37.15 | 8.92 | 10.38 | 12.12 | 21.54 | 21.54 |
| AUC (%) | 97.2 | 96.5 | 96 | 85.5 | 92 | 99.7 | 99.4 | 99.4 | 93.7 | 90.4 |
| Precision | 0.92 | 0.93 | 0.91 | 0.86 | 0.88 | 0.99 | 0.99 | 0.98 | 0.95 | 0.99 |
| Sensitivity (%) | 94.67 | 91.99 | 91.74 | 84.31 | 81.15 | 99.06 | 98.7 | 97.95 | 91.97 | 81.52 |
| Specificity (%) | 92.25 | 93.08 | 91.45 | 86.75 | 88.99 | 98.7 | 98.88 | 98.13 | 95.33 | 99.25 |
| F1 Score | 0.93 | 0.93 | 0.92 | 0.85 | 0.84 | 0.99 | 0.99 | 0.98 | 0.94 | 0.89 |
| 70%-30% | 32 | 64 | 128 | 50 | 16 | 32 | 64 | 128 | 50 | 16 |
| Accuracy (%) | 95.05 | 95.38 | 92.46 | 89.02 | 88.99 | 98.97 | 99.44 | 98.69 | 90.76 | 92.26 |
| Loss (%) | 19.44 | 26.97 | 37.33 | 52.16 | 35.49 | 7.4 | 1.91 | 8.05 | 48.19 | 17.81 |
| AUC (%) | 97.8 | 98.4 | 97.2 | 89 | 93.4 | 99.7 | 100 | 99.6 | 90.8 | 92.3 |
| Precision | 0.96 | 0.96 | 0.95 | 0.9 | 0.9 | 0.99 | 1 | 0.99 | 0.99 | 0.99 |
| Sensitivity (%) | 93.64 | 94.78 | 89.38 | 87.71 | 87.83 | 99.07 | 99.07 | 98.33 | 82.27 | 85.82 |
| Specificity (%) | 96.48 | 95.98 | 95.54 | 90.32 | 90.14 | 98.88 | 99.81 | 99.06 | 99.25 | 98.69 |
| F1 Score | 0.95 | 0.95 | 0.92 | 0.89 | 0.89 | 0.99 | 0.99 | 0.99 | 0.9 | 0.92 |
| 80%-20% | 32 | 64 | 128 | 50 | 16 | 32 | 64 | 128 | 50 | 16 |
| Accuracy (%) | 96.47 | 96.99 | 95.45 | 92.75 | 91.16 | 99.07 | 98.88 | 99.16 | 96.18 | 93.1 |
| Loss (%) | 13.79 | 16.39 | 21.58 | 31.42 | 26.25 | 7.63 | 5.18 | 4.87 | 20.58 | 17.35 |
| AUC (%) | 98.6 | 99 | 98.2 | 92.7 | 97.1 | 99.5 | 99.7 | 99.8 | 96.2 | 93.1 |
| Precision | 0.96 | 0.97 | 0.95 | 0.94 | 0.96 | 0.99 | 0.99 | 0.99 | 0.99 | 0.97 |
| Sensitivity (%) | 96.73 | 97.39 | 95.83 | 90.83 | 86.38 | 99.25 | 99.25 | 99.44 | 92.91 | 88.99 |
| Specificity (%) | 96.2 | 96.6 | 95.07 | 94.65 | 95.94 | 98.88 | 98.52 | 98.89 | 99.44 | 97.2 |
| F1 Score | 0.96 | 0.97 | 0.95 | 0.93 | 0.91 | 0.99 | 0.99 | 0.99 | 0.96 | 0.93 |
| 90%-10% | 32 | 64 | 128 | 50 | 16 | 32 | 64 | 128 | 50 | 16 |
| Accuracy (%) | 98.5 | 98.41 | 96.9 | 96.09 | 92.61 | 100 | 99.63 | 99.63 | 93.38 | 93.84 |
| Loss (%) | 7.27 | 8.75 | 13.84 | 14.8 | 22.28 | 3.66 | 2.57 | 2.5 | 24.7 | 16.41 |
| AUC (%) | 99.4 | 99.6 | 99 | 96.1 | 97.8 | 100 | 99.9 | 100 | 93.4 | 93.8 |
| Precision | 0.98 | 0.98 | 0.96 | 0.95 | 0.9 | 1 | 1 | 1 | 0.89 | 0.98 |
| Sensitivity (%) | 98.69 | 98.37 | 97.5 | 96.88 | 96.23 | 100 | 99.63 | 99.44 | 99.44 | 89.92 |
| Specificity (%) | 98.29 | 98.44 | 96.3 | 95.3 | 88.99 | 100 | 99.63 | 99.81 | 87.31 | 97.76 |
| F1 Score | 0.98 | 0.97 | 0.97 | 0.96 | 0.93 | 1 | 1 | 1 | 0.94 | 0.94 |

**Fig. 5:**
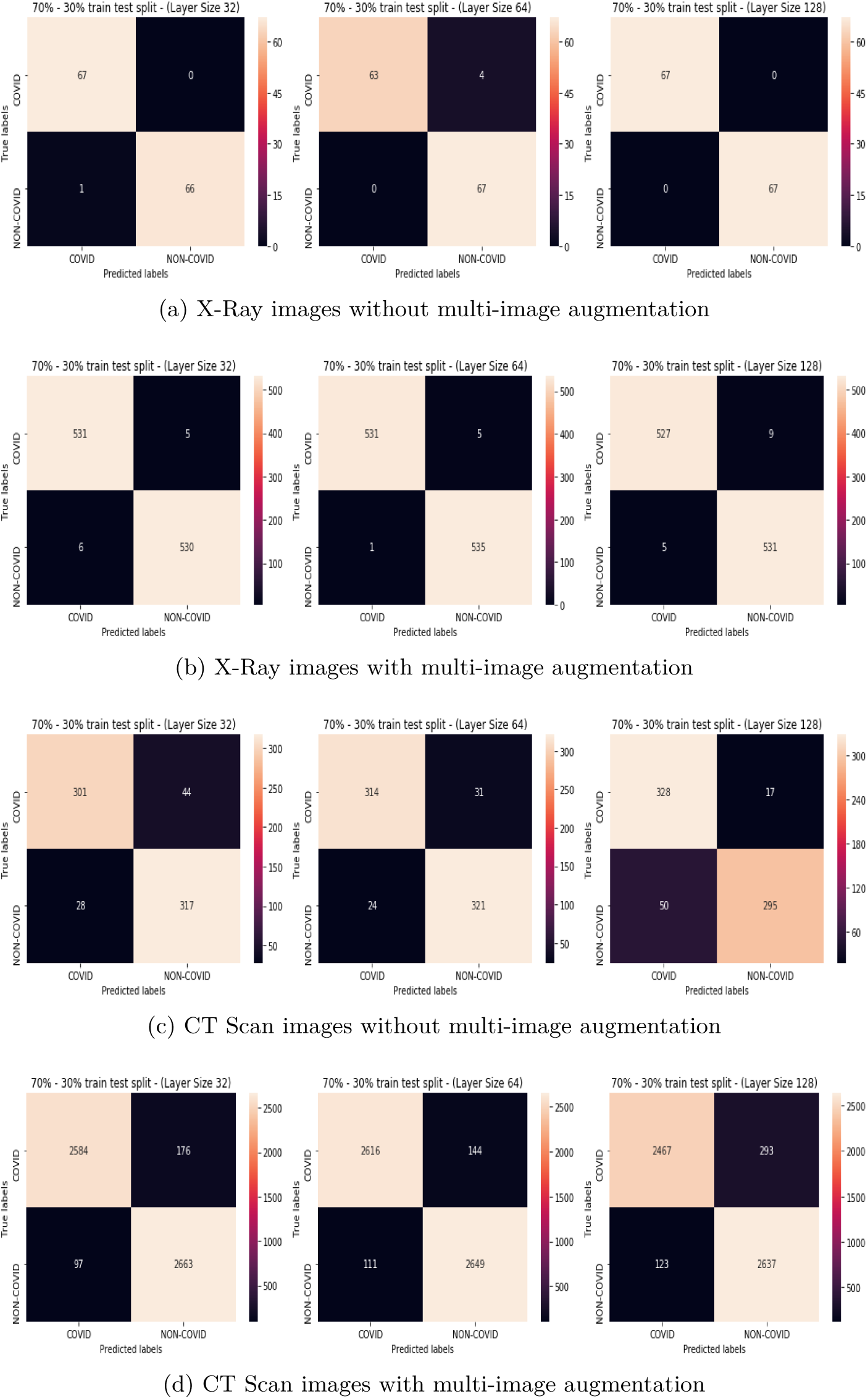
Confusion Matrix of our model when train and test ratio is 70:30

**Fig. 6:**
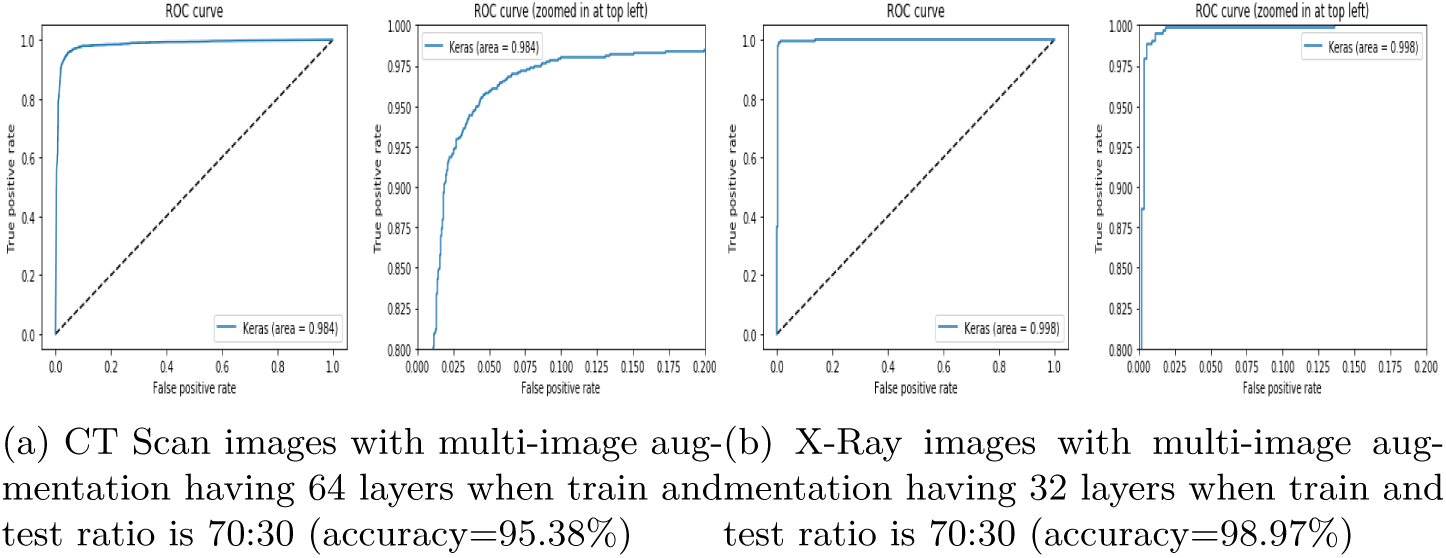
ROC curves

The ROC curve determined on X-Ray images with 64 layers are not shown because it seems like an overfitting having area under the ROC curve as 1. Table 3 shows the COVID-19 screening performance of models having higher accuracy with different train-test values. Figure 7 shows the optimal accuracy comparison between original and augmented techniques on both X-Ray and CT Scan datasets. It is found that in X-Ray dataset, the accuracy slightly differs in both original and augmented techniques due to less number of samples in the dataset. Whereas, in CT Scan dataset, the augmented technique performed with a number of representative images is found better than non-augmented technique with original images. In Table 4 the classification accuracies obtained by the proposed model and the existing models are shown. The existing models either used X-Ray images or CT Scan images for evaluation. Whereas, the proposed augmented deep model has used both CT Scan and X-Ray images of chest with quite large number of samples.

**Table 3:** COVID-19 screening performance of outperforming model with respect to accuracy having different train and test values

|  | Original |  | Augmented |  | Original |  | Augmented |  |
| --- | --- | --- | --- | --- | --- | --- | --- | --- |
|  | CT Scan | X-Ray | CT Scan | X-Ray | CT Scan | X-Ray | CT Scan | X-Ray |
| Train-Test | 60%-40% |  |  |  | 70%-30% |  |  |  |
| Model | Proposed | Proposed | Proposed | Proposed | Proposed | Proposed | Proposed | Proposed |
| Layers | 64 | 64 | 32 | 32 | 64 | 128 | 64 | 64 |
| Accuracy (%) | 90.87 | 99.25 | 93.46 | 98.88 | 92.03 | 100 | 95.38 | 99.44 |
| Loss (%) | 36.13 | 2.59 | 22.82 | 8.92 | 26.81 | 2.22 | 26.97 | 1.91 |
| AUC (%) | 95.2 | 100 | 97.2 | 99.7 | 95.9 | 100 | 98.4 | 100 |
| Precision | 0.9 | 1 | 0.92 | 0.99 | 0.93 | 1 | 0.96 | 1 |
| Sensitivity (%) | 92.17 | 98.51 | 94.67 | 99.06 | 91.01 | 100 | 94.78 | 99.07 |
| Specificity (%) | 89.57 | 100 | 92.25 | 98.7 | 93.04 | 100 | 95.98 | 99.81 |
| F1 Score | 0.91 | 0.99 | 0.93 | 0.99 | 0.92 | 1 | 0.95 | 0.99 |
|  | CT Scan | X-Ray | CT Scan | X-Ray | CT Scan | X-Ray | CT Scan | X-Ray |
| Train-Test | 80%-20% |  |  |  | 90%-10% |  |  |  |
| Model | Proposed | Proposed | Proposed | Proposed | Proposed | Proposed | Proposed | Proposed |
| Layers | 64 | 32 | 64 | 128 | 64 | 64 | 64 | 32 |
| Accuracy (%) | 95.22 | 99.25 | 96.99 | 99.16 | 98.26 | 100 | 98.41 | 100 |
| Loss (%) | 17.16 | 3.84 | 16.39 | 4.87 | 9.47 | 1.08 | 8.75 | 3.66 |
| AUC (%) | 98 | 100 | 99 | 99.8 | 99.5 | 100 | 99.6 | 100 |
| Precision | 0.95 | 0.99 | 0.97 | 0.99 | 0.97 | 1 | 0.98 | 1 |
| Sensitivity (%) | 95.65 | 100 | 97.39 | 99.44 | 99.42 | 100 | 98.37 | 100 |
| Specificity (%) | 94.78 | 98.51 | 96.6 | 98.89 | 97.1 | 100 | 98.44 | 100 |
| F1 Score | 0.95 | 0.99 | 0.97 | 0.99 | 0.98 | 1 | 0.97 | 1 |

**Table 4:**
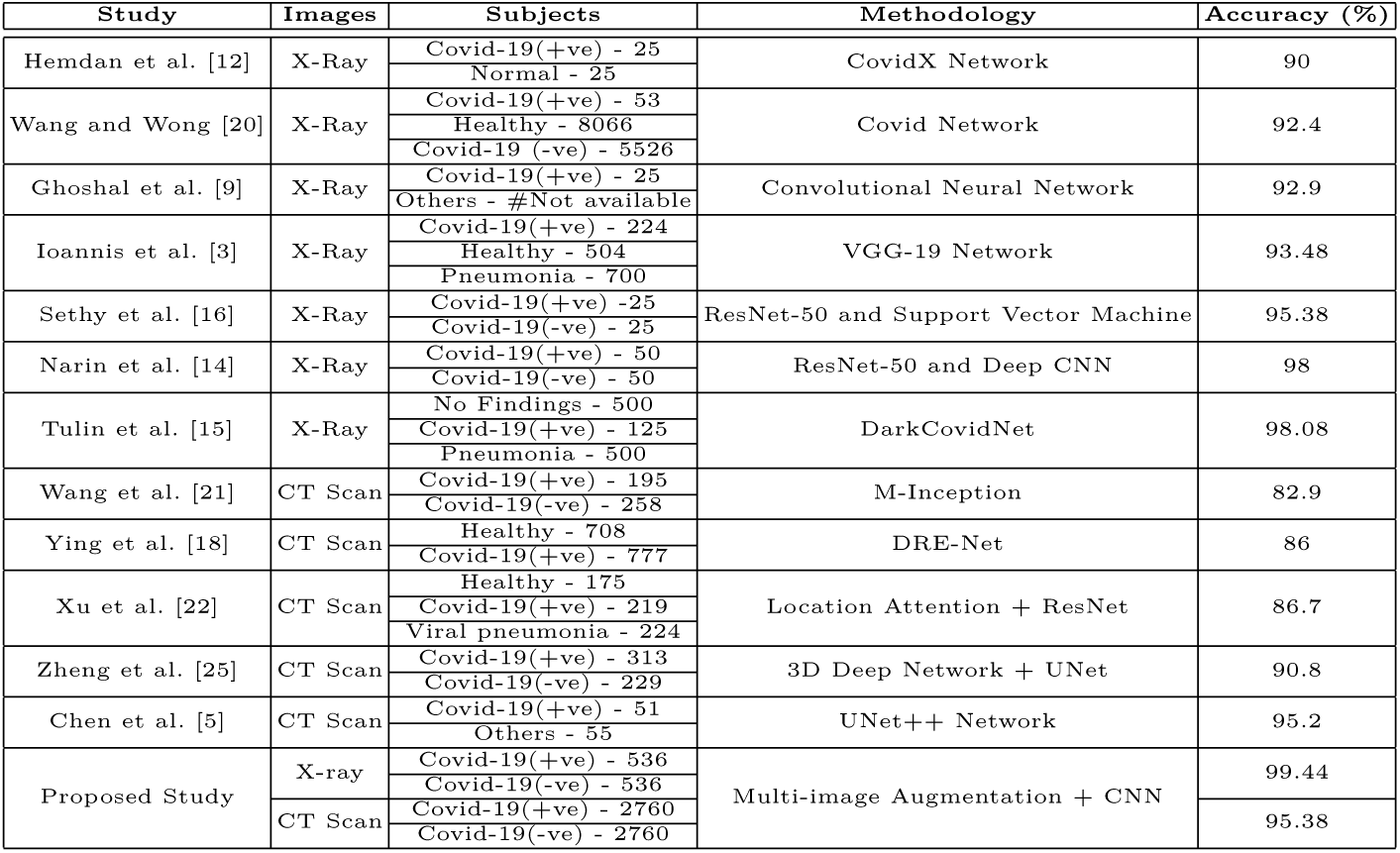
Comparison among various deep learning based COVID-19 screening techniques

**Fig. 7:**
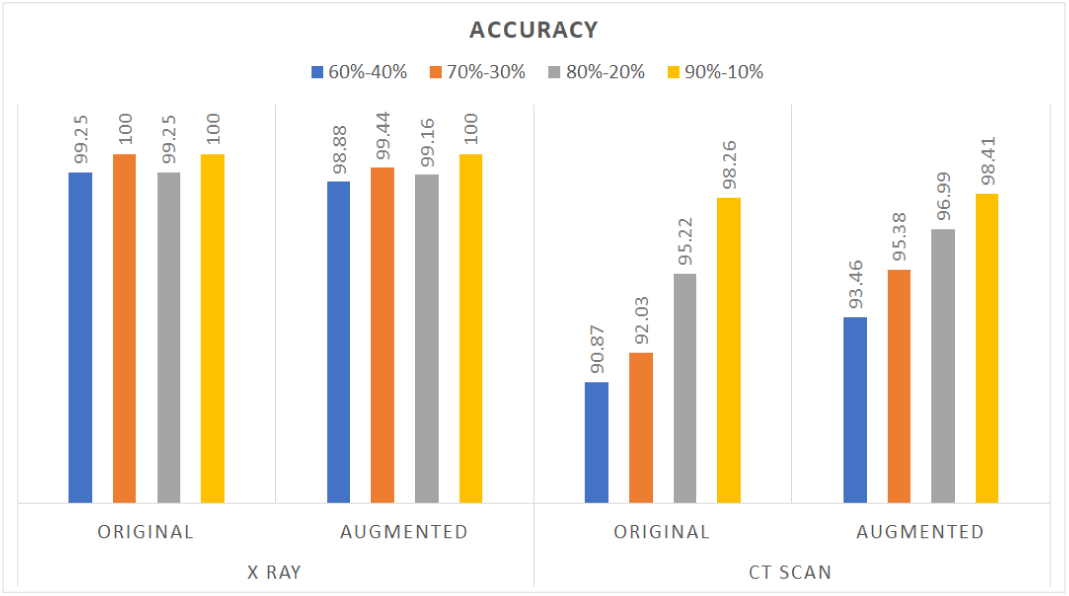
COVID-19 screening accuracy

Hemdan et al. [12] developed a COVIDX-Net for detection of COVID-19 in X-Ray images. An accuracy of 90% is obtained using 25 COVID-19 positive and 25 normal images. Wang and Wong [20] developed COVID-Net which is based on deep neural network for detection of COVID-19. It achieved 92.4% accuracy determined on 53 COVID-19 positive and 8066 normal X-Ray images. Ghoshal et al. [9] used CNN model on 25 COVID-19 positive subjects and obtained the accuracy of 92.9%. Ioannis et al. [3] used transfer learning on 224 COVID-19, 700 pneumonia, and 504 normal images. It obtained 98.75% accuracy for two class and 93.48% accuracy for the three-class problem. Sethy and Behera [16] used CNN models to obtain the image features and classified them by using Support Vector Machine (SVM). They attained 95.38% accuracy using SVM and ResNet50 in combination with 50 images. Narin et al. [14] applied three different deep learning models, i.e., ResNet50, InceptionV3, and Inception-ResNetV2. They achieved 98% accuracy using 50 COVID-19 positive chest X-ray images and 50 normal images. Tulin et al. [15] applied a deep learning model called DarkCovidNet for detection of COVID-19. They used 1125 images which include 125 COVID-19 positive, 500 pneumonia and 500 no-findings to train the model and achieved accuracies of 98.08% and 87.02% for two and three classes classification, respectively.

Wang et al. [21] achieved 82.9% accuracy by applying the modified Inception (M-Inception) deep model using 195 COVID-19 positive and 258 normal CT Scan images. Ying et al. [18] obtained 86% accuracy using 777 COVID-19 positive and 708 normal CT Scan images, with a deep model made on the pre-trained ResNet50, called DRE-Net. Xu et al. [22] achieved 86.7% accuracy for detection of COVID-19 by applying ResNet coupled with 175 Healthy, 219 COVID-19 positive and 224 pneumonia CT Scan images. Zheng et al. [25] introduced a three-dimensional deep CNN model for COVID-19 prediction and obtained 90.8% accuracy using 313 COVID-19 positive and 229 COVID-19 negative CT Scan images. Chen et al. [5] introduced a UNet++ Network for COVID-19 detection and achieved 95.2% accuracy using 51 COVID-19 positive and 55 COVID-19 negative CT Scan images.

Due to less number of X-Ray or CT Scan images that are used for training the deep learning models, often exhibit the classification accuracies not upto the mark. Whereas, the proposed augmented deep model uses large number of X-Ray and CT Scan images for training. This multi-image augmentation has driven the CNN to exhibit higher classification accuracies while a basic deep learning architecture is used. Moreover, the proposed model is compared with ResNet-50 and VGG-16 on the same set of X-Ray and CT Scan images. The proposed augmented deep model outperforms ResNet-50 and VGG-16 as well as the existing models too.

## 4 Conclusion and Future Work

This paper has presented an augmented CNN to detect COVID-19 on chest X-Ray and chest CT Scan images and classify from non-COVID-19 cases. The proposed model need not require to extract the feature manually, it is automated with end-to-end structure. Most of these previous studies have fewer examples for training the deep models. In contrast, the proposed model has used a multi-image augmentation technique driven by first and second order derivative edge operators and this augmentation generates a number of representative edged images. This augmentation technique provides a sufficient number of images to train the model, making the model robust. While, CNN is trained with this augmented images, the classification accuracies of 99.44% for X-Ray images and 95.38% for CT Scan images are obtained. The experimental results are found to be highly convincing and emerged as a useful application for COVID-19 screening on chest X-Ray and CT scan images of corona suspected individuals. The subtle responses of various abnormalities like tuberculosis, pneumonia, influenza and so forth confounds the classifier, restricting the performance of the system. Future works may include detection of multiple conditions such as pneumonia, bronchitis and tuberculosis along with COVID-19 of suspected individuals having respiratory illness.

